# Enabling interpretable machine learning for biological data with reliability scores

**DOI:** 10.1101/2022.02.18.481082

**Authors:** K. D. Ahlquist, Lauren Sugden, Sohini Ramachandran

**Affiliations:** Center for Computational Molecular Biology, Brown University, Providence, RI 02912, USA; Department of Molecular Biology, Cell Biology, and Biochemistry, Brown University, Providence, RI 02912, USA; Department of Mathematics and Computer Science, Duquesne University, Pittsburgh, PA 15282, USA; Department of Ecology, Evolution and Organismal Biology, Brown University, Providence, RI 02912, USA; Data Science Initiative, Brown University, Providence, RI 02912, USA

## Abstract

Machine learning has become an important tool across biological disciplines, allowing researchers to draw conclusions from large datasets, and opening up new opportunities for interpreting complex and heterogeneous biological data. Alongside the rapid growth of machine learning, there have also been growing pains: some models that appear to perform well have later been revealed to rely on features of the data that are artifactual or biased; this feeds into the general criticism that machine learning models are designed to optimize model performance over the creation of new biological insights. A natural question thus arises: how do we develop machine learning models that are inherently interpretable or explainable? In this manuscript, we describe reliability scores, a new concept for scientific machine learning studies that assesses the ability of a classifier to produce a reliable classification for a given instance. We develop a specific implementation of a reliability score, based on our work in Sugden et al. 2018 in which we introduced SWIF(r), a generative classifier for detecting selection in genomic data. We call our implementation the SWIF(r) Reliability Score (SRS), and demonstrate the utility of the SRS when faced with common challenges in machine learning including: 1) an unknown class present in testing data that was not present in training data, 2) systemic mismatch between training and testing data, and 3) instances of testing data that are missing values for some attributes. We explore these applications of the SRS using a range of biological datasets, from agricultural data on seed morphology, to 22 quantitative traits in the UK Biobank, and population genetic simulations and 1000 Genomes Project data. With each of these examples, we demonstrate how interpretability tools for machine learning like the SRS can allow researchers to interrogate their data thoroughly, and to pair their domain-specific knowledge with powerful machine-learning frameworks. We hope that this tool, and the surrounding discussion, will aid researchers in the biological machine learning space as they seek to harness the power of machine learning without sacrificing rigor and biological understanding.

## Introduction

Interpretability has become an increasingly important area of research in machine learning (Azodi, Tang, and Shiu 2020; Freitas 2014), although the term “interpretability” is not always well defined (Lipton 2018; Rudin 2019). When machine learning approaches are applied to biological problems, two major elements of interpretability are desirable: 1) the ability to connect results derived from machine learning with existing biological theory and understanding of biological mechanisms, and 2) the ability to identify and characterize the limitations of a given machine learning algorithm, so that it is applied correctly and appropriately in the biological context of interest. Recent field-specific papers have addressed the first aspect of biological interpretability, by identifying contexts where machine learning approaches are especially powerful, and identifying the types of insights that can be gained (Villoutreix 2021; Gilpin, Huang, and Forger 2020; Zitnik et al. 2019; Lürig et al. 2021; Chen et al. 2018). The second aspect, ensuring that machine learning is applied correctly and appropriately in the biological context, is also a prominent concern in the field, resulting in recent publications suggesting standards for reporting machine learning results in a biological context (Walsh et al. 2021; Wojtusiak 2021; Heil et al. 2021), auditing practices for detecting bias in machine learning applications to biology (Eid et al. 2021), and guides that help prevent common pitfalls in machine learning written for biologists (Jones 2019; Greener et al. 2022).

We previously introduced SWIF(r) (SWeep Inference Framework (controlling for correlation)), a supervised machine learning algorithm that we applied to the problem of identifying genomic sites under selection in population genetic data (Sugden et al. 2018). SWIF(r) learns the individual and joint distributions of attributes (e.g., selection statistics in our motivating application) from training data. SWIF(r)’s algorithm then classifies testing data or application data according to these distributions along with user-provided priors on the relative frequencies of the classes. The use of selection statistics as attributes contributes to the first aspect of interpretability, because the features used for machine learning are derived from existing mathematical theory in population genetics (Weir 1990; Voight et al. 2006; Tajima 1989; Sabeti et al. 2007; Lewontin and Krakauer 1973; Field et al. 2016). Working with a limited number of attributes that are highly meaningful in the biological context, whether summary statistics or other measurements of biological features, creates opportunities for biological interpretation—through the generation of testable hypotheses, or through critical dissection of the classification outcomes.

The generative model underlying SWIF(r) also provides opportunities for interpretability of the second type: characterizing the limitations of the classifier in the context of a specific biological dataset. Here we introduce the concept of a reliability score, a measure of how closely aligned an instance of testing data is to training data, allowing researchers to account for misspecification and missingness in machine learning data. A reliability score ultimately gives users information about the ability of the machine learning model to make a trustworthy classification for a specific instance. The specific reliability score developed here is derived from the Averaged One-Dependence Estimation (AODE) framework underlying SWIF(r). An AODE is a generative classifier, meaning that it learns the attribute distributions (or likelihoods) for each class from training data. Posterior distributions for classes that condition on attributes can then be derived and used to assign class probabilities to new instances. This is in contrast to discriminative classifiers, which learn decision boundaries between classes directly. Because the mathematical form of the reliability score implemented here is dependent on the AODE framework underlying SWIF(r), we refer to the reliability score in this manuscript as the SWIF(r) reliability score, or SRS. While Sugden et al. (2018) demonstrated SWIF(r) to be reasonably robust to misspecification of training data and missing or undefined statistics in the context of selective sweeps, the SWIF(r) probabilities reported do not offer insight into the degree of “trustworthiness” of a particular classified instance. Other classification approaches using machine learning, including Logistic Regression classifiers, k-Nearest Neighbors classifiers, Support Vector Machines, Neural Networks and Bayesian classifiers suffer from the same limitation: that is, users will receive classification outcomes from their models, with little visibility into the underlying features contributing to classification (Murphy 2012; Rudin et al. 2022).

The SRS is of particular value for biological applications where it is necessary to use simulated training data, a practice that is ubiquitous in population genetics, and extends to many evolutionary questions across different areas of biology (Villanea and Schraiber 2018; Sankararaman et al. 2014; Schrider and Kern 2016; Pavlidis, Jensen, and Stephan 2010; Battey, Ralph, and Kern 2020b, [a] 2020). Simulation studies also underlie other analyses of complex biological phenomena, such as cancer progression (Han et al. 2019; Zangooei and Habibi 2017). In many cases, researchers evaluate the performance of an algorithm or method on simulated “testing” data, before applying it to “application” data drawn from biological samples. In order for these models to be reliable, there needs to be a high degree of similarity between simulated data and application data. Despite best efforts, researchers must often contend with differences between simulated and application data, arising from various sources. When these differences arise, it may be preferable to set aside data instances with poor fit, allowing the classifier to effectively abstain when it encounters signatures it has not been trained to expect; one goal of the SRS is to enable the user to identify such poor-fit instances. For example, in the context of genome scans for selection, such unexpected signatures can arise in regions of the genome with unusual statistical signatures due to idiosyncratic recombination or mutation profiles that are not captured in simulated training data, such as the MHC (de Bakker et al. 2006; Lenz et al. 2016).

In this study, we assess the ability of the SRS to identify deficiencies affecting the classifications produced by SWIF(r) across different biological applications and contexts. Although SWIF(r) does return probabilities to facilitate interpretation of its results, like any machine learning classifier, SWIF(r) is restricted by its design, including the choice of training data and specified classes under consideration. When a classifier encounters data that is not well-matched to any trained class, its behavior can be erratic, and at worst, deceptive, as we illustrate in our experiments. In particular, we study performance in situations in which any of the following occur: 1) classes of data are missing in the training data but present in the application data, 2) there exists a systemic mismatch between training data and application data, 3) there are missing attributes for valuable instances of application data.

## Results

In this study we introduce the SWIF(r) Reliability Score (SRS), and explore its utility for applications to biological data. We define the SRS, and use simulations to demonstrate the sensitivity of the SRS to underlying data features, including the distribution of individual attributes and the structure of correlation between attributes (Figure 1). Next, we look at a series of biological datasets to illustrate the utility of the SRS in applications to biological problems. First, we use a wheat morphology dataset to study the effect of a class that is present in testing data but unaccounted for in training data (Figure 2). Second, we use phenotype data drawn from the UK Biobank to study the effect of a systemic mismatch between training data and testing data, in this case due to differences in ancestry or sex across the training and testing sets (Figure 3). Third, we use simulated data and 1000 Genomes Project data to illustrate the effects of missing data on classification performance in a population genetics context (Figure 4).

**Figure 1.**
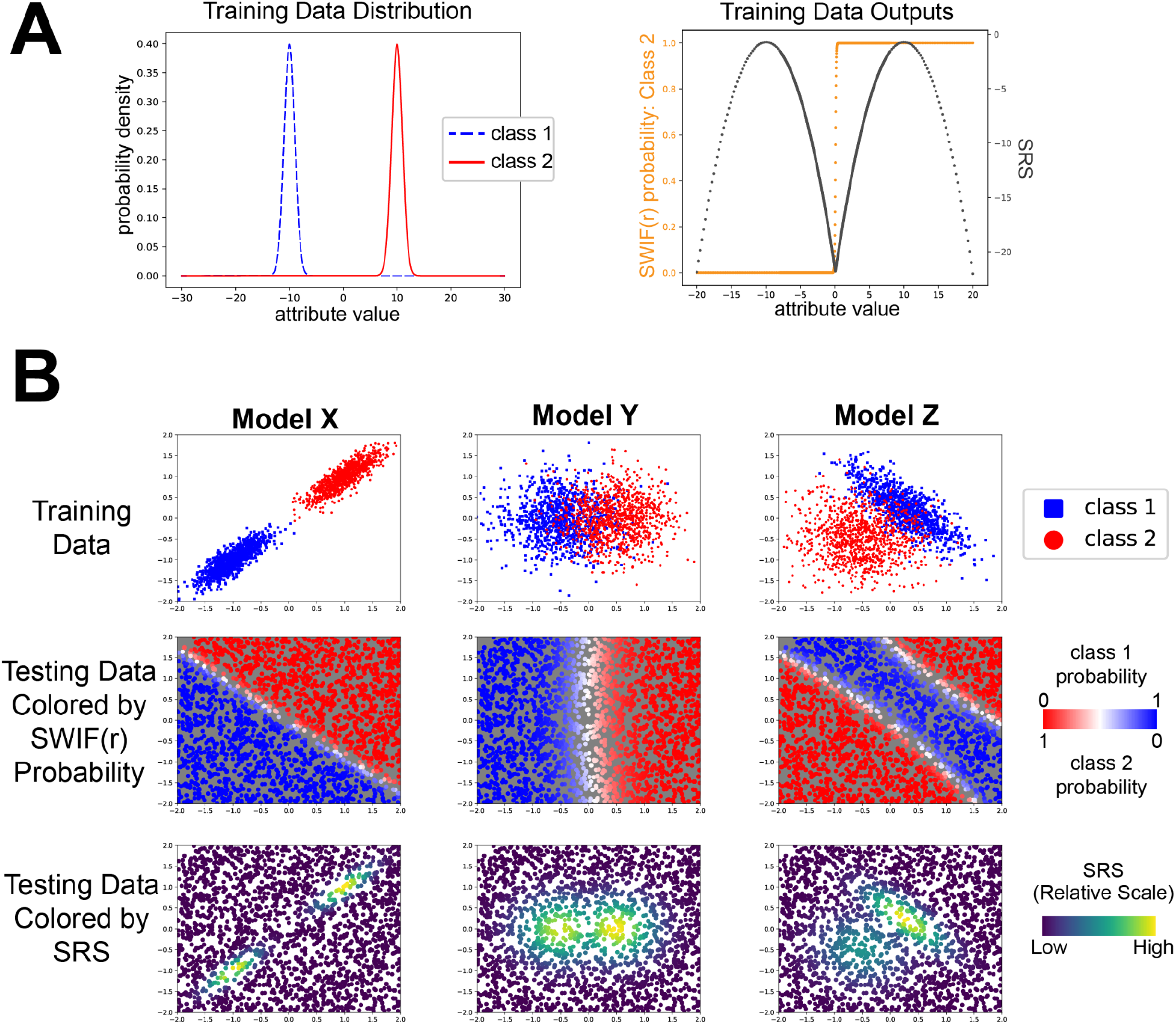
The SRS reveals the underlying structure of the data in example training scenarios. **A)** For a classification problem with a single attribute, we illustrate how the SRS rises and falls over a range of classification probabilities. Data was generated for two classes (class 1: blue, dashed line, and class 2: red, solid line) with a single attribute. SWIF(r) was trained using 1000 instances of each class drawn from the attribute distribution shown on the left. Following training, SWIF(r) was tested on values across the range (−20, 20), resulting in the SWIF(r) probabilities and SRS values shown on the right. In the right hand graph, SWIF(r) probability of class 2 is shown in orange, and SRS is shown in dark gray. Here we see that the SRS drops in regions that are not represented by either class in training, across a wide range of classification probabilities. **B)** SRS detects differences in the correlation structure of testing data compared to training data in a two-attribute model. Data was generated for two classes (class 1 shown in blue, and class 2 shown in red) with two attributes. Different distributions and correlations between attributes were chosen to create three SWIF(r) models labeled “X” (left), “Y” (middle) and “Z” (right). In the top row for each model is the data used to train the model, with one attribute along each axis. The middle and bottom rows show testing data drawn uniformly across the same area shown. In the middle row, testing data is colored by the SWIF(r) probability, ranging from 0 to 1 for each class (note that for a binary classifier, *P*(*class* 1) + *P*(*class* 2) = 1). In the bottom row testing data is colored by the relative value of the SRS for the testing data shown, with yellow corresponding to the highest SRS, and purple corresponding to the lowest SRS.

**Figure 2.**
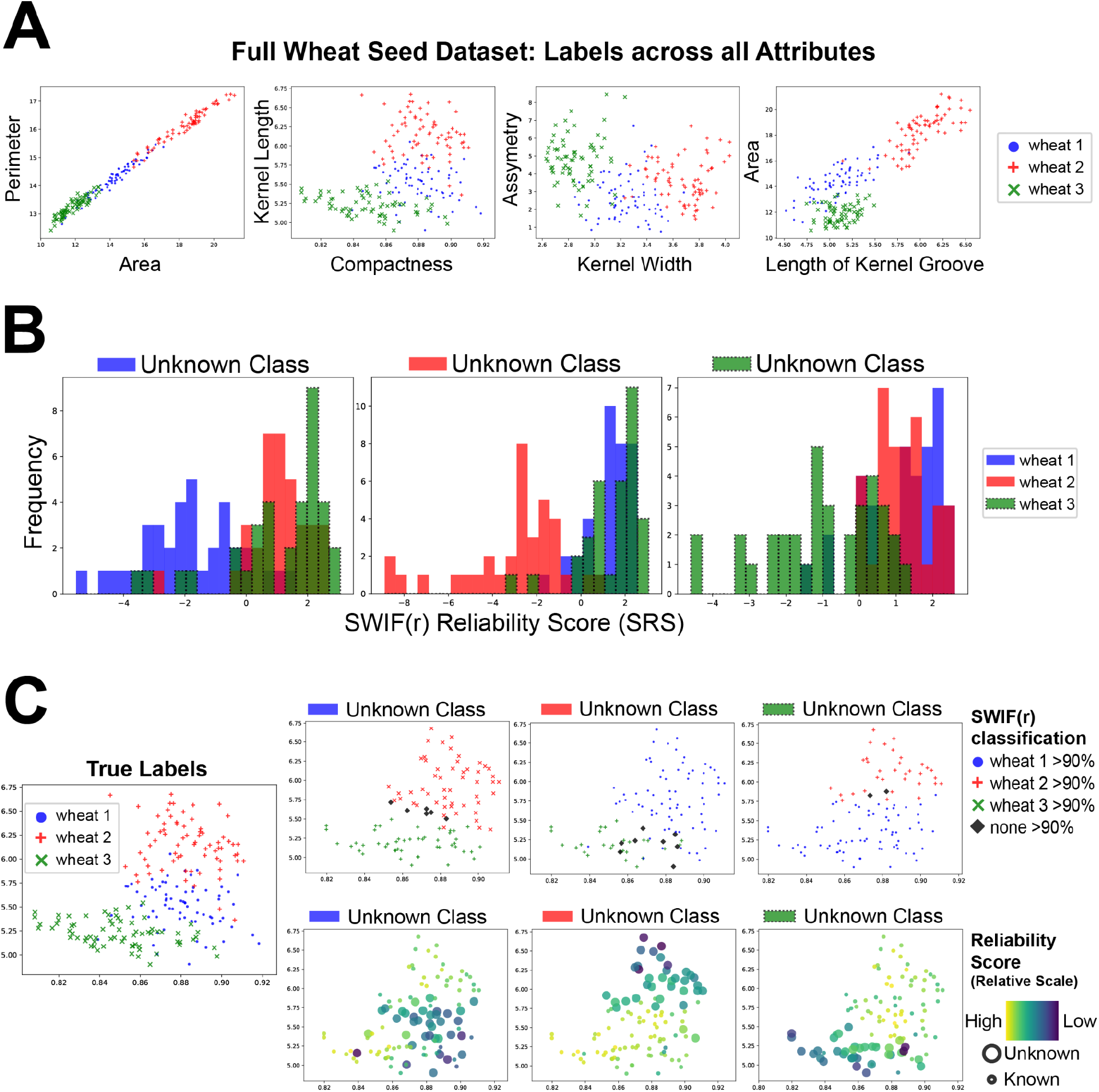
The SRS is lower when an instance’s class is excluded from training. SWIF(r) was trained using each combination of two classes in the wheat morphology dataset (see Methods, (Charytanowicz et al. 2010)). The trained model was then tested on instances from all three classes. **A)** Distribution of the training and testing data across all seven attributes: Perimeter, Area, Kernel Length, Compactness, Kernel Width, Asymmetry and Length of Kernel Groove. Note that wheat 1 (blue circles) generally has trait values that are intermediate to the other two classes **B)** Histograms show the distribution of the SRS for instances of each class. In each case, the unknown class has a more negative distribution of SRS compared to the two known classes. This is true even for wheat 1 (blue), which has attributes that are intermediate in value as compared to the other classes. **C)** Data graphed by two attributes (Kernel Length and Compactness). Left: full data set colored by true labels. Right: colored by SWIF(r) probability (top) and SRS (bottom). SWIF(r) scores were generally greater than 90% for one of the two trained classes, *even for instances from the unknown class*, with just a handful of points receiving intermediate values from SWIF(r) (black diamonds). In contrast, coloring by SRS shows that points associated with the unknown class (larger dots) tend to have lower SRS, while points associated with known classes (smaller dots) received higher SRS.

**Figure 3.**
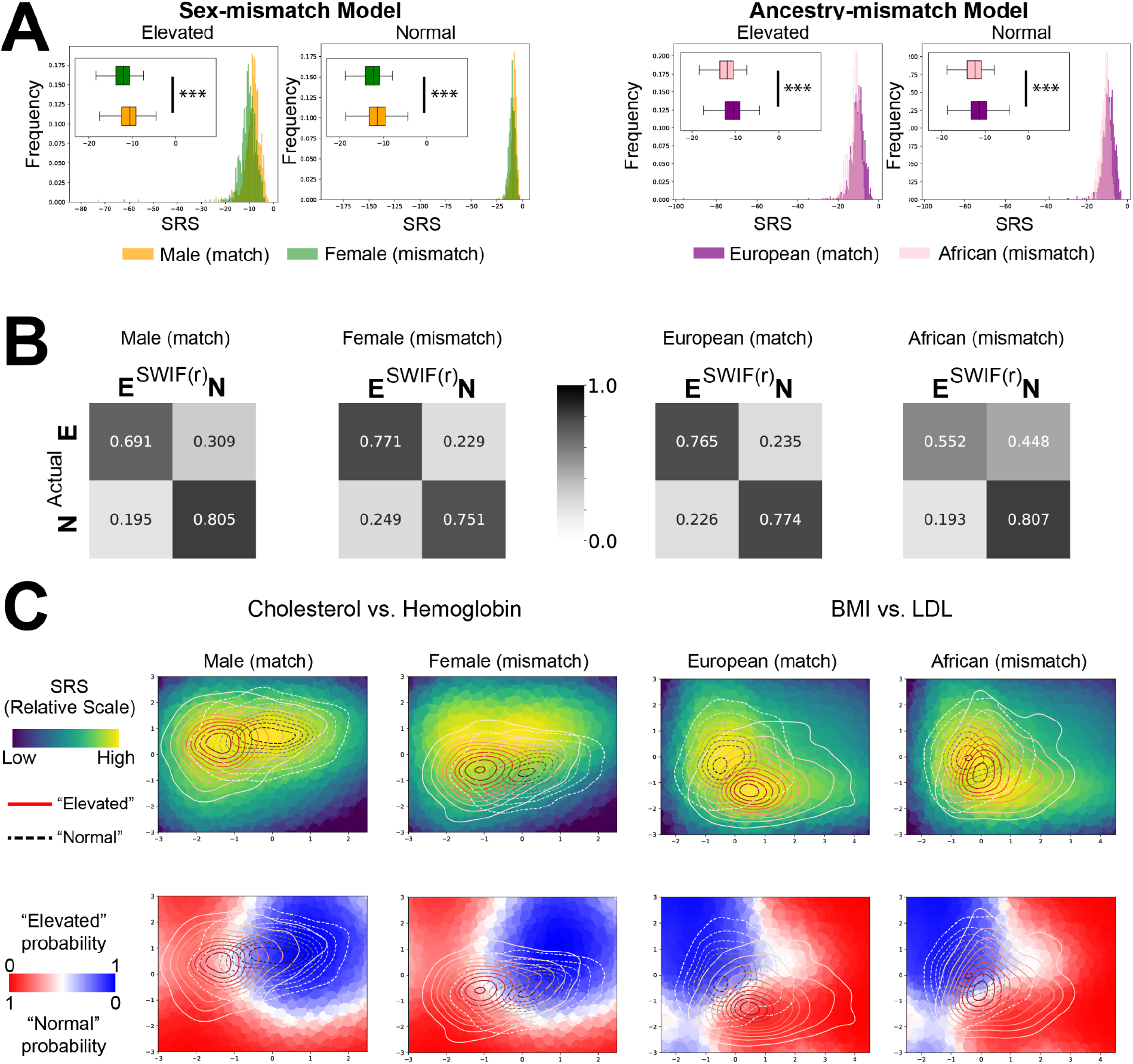
Low average SRSs can indicate systemic mismatch between training and testing data. **A)** In the sex-mismatch model (left), SWIF(r) was trained using a dataset consisting of male individuals of European ancestry divided into two categories based on HBA1C readings, used to esimate blood sugar: Elevated, or Normal. The data provided for learning consisted of 22 health-related attributes (see Methods). The trained model was then tested on two cohorts, females with European ancestry and an independent cohort of males with European ancestry. These cohorts were labeled with their Elevated or Normal status, allowing for identification of correct or incorrect classification by SWIF(r). In the top left, we see the distribution of SRS for Elevated and Normal individuals from either the matching cohort (male) or non-matching cohort (female). The female cohort has lower average SRS, visible as a leftward shift in both the Elevated (p= 1.53e-4) and Normal (p= 5.11e-34) distributions (*** represents p<0.001). Likewise in the ancestry mismatch model (right), SWIF(r) was trained using a dataset of male and female individuals of European ancestry divided into two categories: Elevated and Normal. The trained model was then tested on two cohorts, males and females with African ancestry and an independent cohort of males and females with European ancestry. As above, we see the distribution of SRS for each cohort. The non-matching African ancestry cohort has lower average SRS, visible as a leftward shift in both the Elevated (p= 6.04e-06) and Normal (p= 1.50e-28) distributions. **B)** Confusion matrices show differences in SWIF(r) classification accuracy between matching and non-matching cohorts. The non-matching sex cohort experienced a small shift towards the Elevated classification when compared to the matching cohort. The non-matching ancestry cohort experienced a larger shift towards the Normal classification when compared to the matching cohort. **C)** SRS and SWIF(r) probability are calculated over a plane defined by two of the twenty-two model attributes, providing a background of points for each graph. On top of each is graphed a contour plot of the distribution of Elevated (red-to-white, solid line) or Normal (black-to-white, dashed line) data for each cohort. On the left, comparing the Male and Female cohorts we can observe a shift in the overall mean of the data, pushing the Female cohort into an area with lower SRS values (top) and greater Elevated SWIF(r) probability (bottom). On the right, comparing the European and African cohorts, we observe that the mean difference between the Elevated and Normal cohorts is higher for individuals with European ancestry, and smaller for individuals with African ancestry. This results in greater overlap between the Elevated and Normal distributions in the African cohort, as well as an overall shift towards the Normal classification.

**Figure 4.**
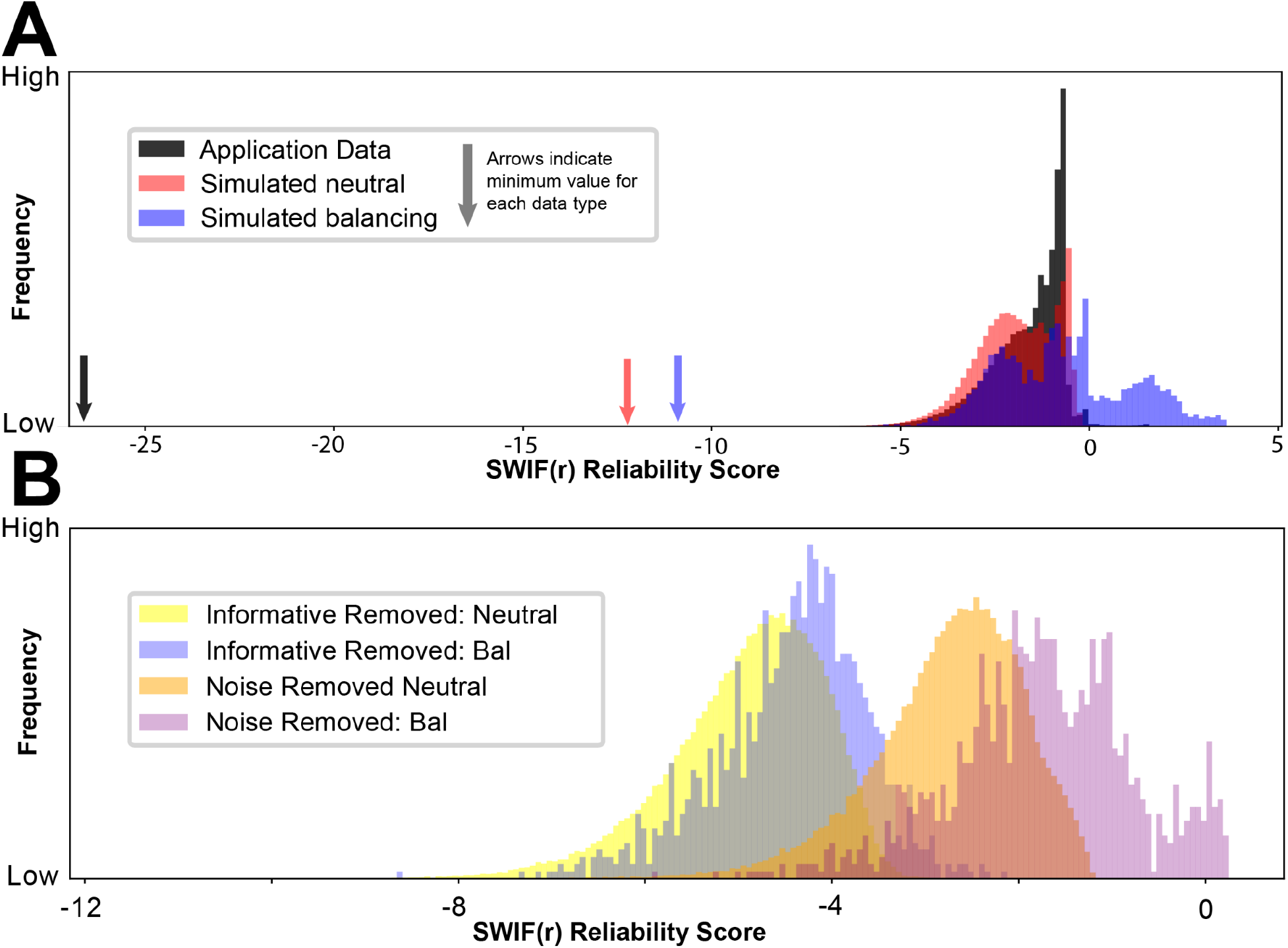
The SRS can identify valuable attributes to create smarter filters on missingness. **A)** Application data exhibits extreme low values of the SRS as compared to simulated data. **B)** Missing Informative attributes lead to lower SRS compared to missing Noise attributes. SWIF(r) was trained with a combination of five potentially informative attributes (“Informative”), and three “Noise” attributes drawn randomly from Gaussian distributions with no relation to the class of the data. In the testing set, 3 attributes were removed from each instance: either all three Noise attributes (“Noise Removed”), or 3 randomly selected Informative attributes (“Informative Removed”). The SRS was then calculated for all instances. Instances where Informative attributes were removed have lower average SRS values compared to instances where the same number of Noise attributes were removed, visible in the histograms shown.

### Overview and Definition of SRS

The SRS, developed here for SWIF(r) but generalizable to other generative classifiers, is an interpretability aid built from intermediate steps within SWIF(r)’s classification pipeline. SWIF(r) classifies instances using Averaged One-Dependence Estimation, an extension to Naive Bayes that incorporates pairwise joint distributions of attributes to estimate the joint likelihood of the observed data conditioned on each class:

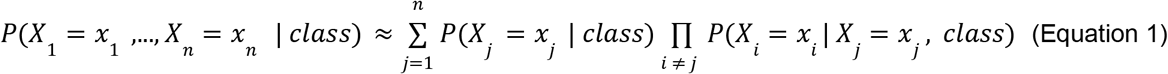

Where X_i_ represents the i^th^ attribute and n is the number of attributes. With the use of prior closs probabilities, and a denominator representing the marginal probability of the data over all classes, SWIF(r) returns a posterior probability for each class (see Materials and Methods, Equation 3). The SRS focuses solely on the estimated likelihoods, because these are the values that indicate the fit of the instance to the trained model, while the posterior probabilities convey the *relative* fit across the trained classes. Because the posterior probabilities are forced to sum to 1, instances that do not resemble any trained class may still receive a high classification probability for the “least bad” option when classified by SWIF(r); the SRS is designed to help detect such instances.

To calculate the SRS, we calculate the estimated joint likelihoods, taking the maximum across all classes, then apply a log_10_ transformation to aid visualization:

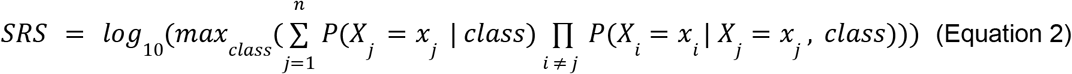

By taking the maximum with respect to class, the SRS is agnostic to the specific class labels of an instance. Because the SRS depends heavily on the specifics of a particular application, SRS values cannot be compared across experiments. Instead, SRS are designed to be compared *within the context of a single trained model*.

In Figure 1, we illustrate how our SRS works in the context of two examples. In the first example (Figure 1B), classification into one of two classes is based on a single attribute, essentially reducing the classification problem to a straightforward application of Bayes’ rule. Observations in class A follow a Gaussian distribution with a mean of -1 and standard deviation 0.2, while observations in class B follow a Gaussian distribution with a mean of +1 and standard deviation 0.2. Note that in this case, the SRS reduces to the maximum value of the two probability density functions. As we classify values across the x-axis, we see that the probability assigned to class B monotonically increases, as we would expect. However, the SRS dips in three places: values to the left of class A, values between the two classes, and values to the right of class B. In these ranges, we have values of testing data that are a poor match to both classes. We note in particular that low SRS values can occur across the full range of classification probabilities, including probabilities that would be interpreted as “very likely A” to “very likely B” and everything in between. This makes the SRS a valuable, complementary addition to the standard classification output.

In Figure 1B, we scale up to examining a series of models with two attributes. The distribution of the training data is shown in the first row of panel B. The second row of panel B shows the SRS evaluated at points drawn uniformly across the plane. We see that SRS is highest when testing points are similar to training data, not only in attribute value, but also with respect to the correlation between attributes. In particular, we see that for instances in which individual attribute values are all within typical ranges, the SRS will still decrease if they jointly lie in a region of space not well-represented in the training data. This illustrates the sensitivity of the SRS to the high-dimensional structure of the trained dataset, which allows it to detect “unexpected” data even when no single attribute value constitutes an outlier with respect to its trained distribution. The bottom row shows the SWIF(r) probability evaluated at points drawn uniformly across the plane. Since SWIF(r) tries to find the “best” class to fit a particular instance, we see regions of the plane in which instances lie far from the training data of both classes, as captured by the SRS, yet are classified with high probability to one class or the other. More generally, this acts as an illustration of the inherent limitations of a classifier as a set of decision boundaries. Combining standard classification output with the SRS enables a more complete and nuanced interpretation of results.

### The SRS is sensitive to instances from unknown classes

The SRS is sensitive to instances from classes in application data that are not present in the training data. While SWIF(r) is forced to find the best match across trained classes, using the SRS allows researchers to identify those instances that are unusual with respect to both classes (Figure 1), and, potentially, to associate them with a class that was excluded from training or was previously unknown. In order to demonstrate the effect of missing classes in the training set in an authentic biological context, we analyzed a dataset containing seven morphological attributes of seeds from three types of wheat. By considering the three types of wheat as three classes, this gives us an opportunity to investigate the performance of SWIF(r) and the SRS in the case where one of the three classes is unrepresented in training. To this end, we trained three SWIF(r) models, each one excluding one of the three classes. We then applied SWIF(r) to the full testing set, which contained instances from all three classes. Figure 2 shows that for each model, the instances belonging to the class that was excluded from training have dramatically lower SRS values on average than instances that belong to a known class. This is even true when the unknown class contains values that are intermediate to the two known classes, suggesting that the SRS is able to indicate deficiencies in the training set without relying on the presence of extreme values.

### The SRS is sensitive to systemic bias in training data relative to application data

In Figure 3, we demonstrate how the SRS can be used to identify systemic differences between training data and application data, by examining examples of demographic mismatch between training and testing sets in the UK Biobank. Two models were trained to predict elevated or normal blood sugar levels (HBA1C>48 mmol/mol for “Elevated”, HBA1C<42 mmol/mol for “Normal”) from a collection of phenotypes as attributes. In the first “sex mismatch” model (3A, left), the model was trained on data from males with European ancestry, using 22 attributes (see Methods) to predict whether individual blood sugar would be Elevated or Normal. This model was then tested on a second cohort of males with European ancestry, and a “non-matching” cohort consisting of females with European ancestry. In the second “ancestry mismatch” model (3A, right), the model was trained on data from male and female individuals with European ancestry, using the same 22 attributes, with the same classification goal. This model was then tested on a second cohort of male and female individuals with European ancestry, and a non-matching cohort consisting of male and female individuals with African Ancestry (see Methods). In both cases, the average SRS of the non-matching cohort was lower than the average SRS of the matching cohort (Figure 3A, Table S1). This corresponded with differences in the way that SWIF(r) classified the non-matching cohort (Figure 3B). Compared to the matching cohort for each experiment, in the non-matching cohorts, SWIF(r) became systematically biased toward one of the two classifications, classifying more individuals as Normal in the African ancestry cohorts, and more individuals as Elevated in the female sex cohorts (Figure 3B).

In order to investigate the causes of the drop in SRS, we identified some attributes with large mean differences between matching and non-matching cohorts. In Figure 3C left, we show a pronounced example of mean difference between the distributions of cholesterol and hemoglobin across male and female cohorts that results in lower SRS values for the female cohort. In the background of the upper plots, we plot the value of the SRS across a plane defined by those two attributes, and overlay the distribution of trait values for each cohort. In the lower panels of Figure 3C, the background is colored using SWIF(r) probability. This allows us to visualize how the attribute values occupy different ranges of values of the SRS and of SWIF(r) probability. From this view, we can notice some patterns emerging in this mean shift example. The matching (male) cohort sits in the center of the area with highest SRS. Likewise, the probability distributions determined using the (male-only) training data appear to do a good job separating the testing data from the male cohort. In contrast, the female cohort is shifted downward, out of the area of the plane with the highest SRS. This also causes the probability distribution shift relative to the means of this cohort, resulting in a greater number of individuals from both Elevated and Normal classes being classified as having elevated blood sugar levels. In the ancestry mismatch example, we observe that there is a smaller mean difference between the classes of the mismatch ancestry cohort (African/elevated vs. African/normal) than there is between the matching ancestry cohort (European/elevated vs European/normal). This demonstrates a major failure for the classifier: by relying on a difference that is present in the training cohort, the classifier will not be able to accurately distinguish between individuals belonging to a cohort that lacks this difference. This results in the increased classification of individuals with African ancestry as Normal, for individuals in both the Normal and Elevated categories. We can see this in the lower right panel, where the distribution of individuals with elevated blood sugar levels is shifted towards the area given a high Normal probability (blue) or intermediate probability (white), rather than the area with high Elevated probability (red). The SRS lacks sensitivity in this scenario: *if differences between cohorts cause individuals from one class to look like valid entries from another class, SRS cannot help to identify these misclassified individuals*. This represents a situation under which we would expect all classifiers to fail.

### The SRS can be used to filter instances missing valuable attributes

We have shown how SRS can assist researchers in detecting unknown classes in testing data (Figure 2), and identifying systemic mismatches between training and testing data (Figure 3). Another feature of SRS that is especially relevant to real-world data is how it responds to data ‘missingness’. Many machine learning classifiers do not allow for any attributes to be missing from an instance for classification to be performed. SWIF(r) allows for any number of missing attributes, as long as at least a single attribute is defined. However, while SWIF(r) is tolerant of missing data, performance generally declines as missingness increases (Sugden et al. 2018). Knowing this, it may be tempting to set a simple threshold on the number of missing attributes allowed, or otherwise filter instances that contain missing attributes.However, we find that the SRS offers a method for selectively filtering instances that are missing valuable attributes, without losing instances that can be classified confidently with existing attributes.

In Figure 4, we illustrate this in a population genetics context, in which SWIF(r) is trained on simulated training data representing neutral evolution and balancing selection, with application data coming from the 1000 Genomes project (1000 Genomes Project Consortium et al. 2015, Materials and Methods). In this dataset, each instance is a single site in the genome. Missing data is ubiquitous in both simulated and application data due to the constraints on calculating many of the attributes, which in this case are selection statistics. In Figure 4A we observe the general problem that low SRSs are present, especially in application data. As we know from prior experiments, this can have multiple causes related to mismatch between training and testing data. In this case, application data had the highest average number of missing attributes, missing an average of 2.87 attributes out of five total attributes. Balancing selection simulations were missing an average of 2.75 attributes, and neutral evolution simulations were missing an average of 1.56 attributes. The pattern of missing attributes differed across data types and SRS ranges. Notably, out of 23 application data instances with SRS values less than -16, all 23 were missing iHS and nSL, and 22 out of 23 were missing DDAF.

In Figure 4B, we set up an experiment that demonstrates the benefit of using the SRS to determine how to filter instances with missing data, rather than setting an arbitrary threshold on missingness. First, we created a filtered training dataset from simulated data, using only instances for which all of the calculated statistics are present. We label our original set of selection statistics “Informative attributes”. We also added 3 additional attributes that have no connection to instance properties, instead they are drawn randomly from Gaussian distributions. These “Noise attributes” are drawn in an identical manner for both training classes, so they have no information value to the machine learning algorithm. We filtered testing data in the same manner, with the same added Noise attributes. As a final step in the creation of testing data for this experiment, we added missingness by eliminating either three Informative attributes (Figure 4B, yellow and blue) or the three Noise attributes (Figure 4B, orange and red). Holding the number of missing attributes constant, we see that SRS is lower when Informative attributes are missing, and higher when Noise attributes are missing. A strict threshold on missingness would treat Noise and Informative attributes equally when deciding to eliminate an instance from consideration by the classifier. Instead, we demonstrate that we can use SRS to identify valuable attributes, in order to avoid eliminating instances that are missing low-value or uninformative attributes.

## Discussion

In this study we introduce a new concept for scientific machine learning studies, the reliability score, designed to provide information about how closely a given testing instance matches with the model of the data underlying a classifier. We develop a version of the reliability score, the SWIF(r) Reliability Score or SRS, and illustrate the utility of the reliability score using multiple biological datasets. Current debates in the machine learning field propose two major competing objectives for developing machine learning frameworks: interpretable methods, which have inherent features which allow users to understand how model conclusions are reached (Rudin 2019; Rudin et al. 2022), and explainable methods (and/or explanatory tools) which provide post-hoc or mathematically distinct overlays onto model conclusions in order to aid human understanding of how model conclusions may have been reached (Linardatos, Papastefanopoulos, and Kotsiantis 2020). In this paper, we land in the first camp: using features of SWIF(r)’s underlying algorithm to draw scrutiny to multiple failure cases, we can leverage SWIF(r)’s inherently interpretable, generative framework to build the SRS. The SRS allows a user to probe for important deficiencies in the training and testing data, including missing classes, systemic mismatch between training and testing data, and instances missing informative attributes. Understanding the causes that contribute to a low SRS offers researchers several potential remedies, including adding more informative attributes to their analysis, expanding or revising the training set or excluding instances with low SRS from their analysis.

The SRS represents a valuable measure for SWIF(r), and similar reliability scores could be created for closely related methods with a generative basis. This underscores the inherent interpretability of generative methods: because they learn the underlying distribution of data for each class, we can reveal elements of that process in order to improve the quality of classification, and understand it’s basis. Since discriminative classifiers learn decision boundaries directly without learning underlying distributions, these methods (including most neural networks, support vector machines, random forest classifiers, regression-based methods and more) will require different means of measurement to get at similar cases of classification failure. It is possible to use the SRS independent of applying SWIF(r), as a complementary read-out alongside another machine learning classifier. In this case, the SRS would be an explanatory tool for examining results or refining the design of the training set, but could not be understood as providing insight into the specific classifier. When used independently of SWIF(r), the SRS effectively provides a form of outlier detection. One major benefit of using the SRS versus a simpler form of outlier detection is that the SRS uses the structure and correlations of the training data, rather than just extreme values to identify instances that are unexpected or unusual. This is especially well demonstrated in Figure 2, where we can use the SRS to identify instances of the wheat 1 class, despite the attribute values of the class being intermediate between the other classes. If a similar method was developed solely for the purpose of providing an explanatory method, it could potentially be improved by considering higher-dimensional correlations, or by testing different simplified models of the data distributions, instead of using the data model underlying SWIF(r).

One limitation of the SRS is that while the SRS indicates problems with missing data or deficiencies in the dataset, a solution to those problems may not always be possible or simple to implement. Researchers will need to rely on domain specific knowledge to decide whether the indicated problem can be solved by adding additional classes or attributes to their model, or by including more, or more diverse instances, in their training data. Discovering a solution to low-reliability instances will require answers to subjective and context-dependent questions that may be difficult to anticipate and may not generalize. For example, researchers may face arbitrary decisions about the number and types of classes to include in training and cost-benefit considerations when deciding whether to gather the additional data required to add new attributes or additional training examples. How a researcher goes about designing their analysis will influence their results, and important questions remain about how to set standards and guide these decisions. Increasing transparency and publication of negative results could help address some of the problems raised by these subjective questions.

This paper comes at a time when the investigation of flaws in machine learning has become an important topic of research in its own right. As machine learning methods are put to the test for applications like medical image analysis, we encounter examples where the accuracy and value of machine learning methods are artificially inflated by flaws in the training set, overconfidence in results, or reliance on unintended features of data (Poplin et al. 2018; Badgeley et al. 2019). Many of these machine learning methods are so-called ‘black box’ methods that are relatively difficult to deconstruct. Explanatory tools built to analyze the output of these methods often rely on post-hoc challenges to a trained model. In contrast, the relative mathematical simplicity of SWIF(r) and the SRS allows for more direct observation.

Across many areas of biology there is an opportunity to improve machine learning analyses by creating community standards and best practices for simulation, interpretability and reproducibility. As machine learning applications to biological data expand, it will be important to prioritize the ability to draw biological conclusions from data. We also can not afford to lose sight of the way that subjective decisions guide the design of machine learning models. Domain-specific biological knowledge currently provides the best available guide for navigating the complex, multi-factorial choices involved in model design. In order to best leverage this domain-specific expertise, we need to ensure that machine learning models are easy to interpret, dissect and critique, even by researchers without machine learning backgrounds. Tools that enhance the interpretability and transparency of machine learning algorithms are an important step towards this goal.

## Materials and Methods

### Overview and Definition of SRS

SWIF(r) was originally defined using the following formula:

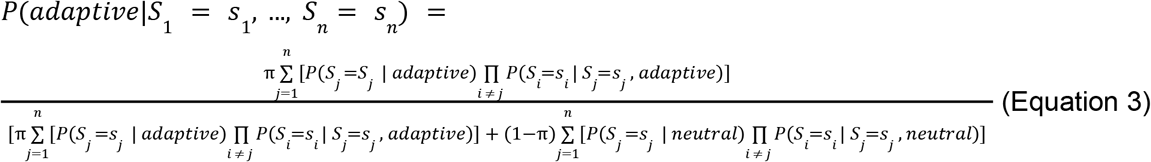

This framework can be generalized to other applications by simply changing the classification categories (in Sugden et al. (2018), adaptive and neutral), and with adjustment of the Bayesian prior *π* according to the specific application. Attributes, denoted here as “S” for Summary Statistic, must be continuous variables. Equation 1 (Results) adapts the numerator of SWIF(r), describing how the classifier learns the likelihood of the observed data conditioned on each class using both the marginal distributions of the attributes and the joint distributions of the attributes, from which conditional distributions are derived. The SWIF(r) probability is the posterior probability of a class given the set of attributes. The output from Equation 3 can be thought of as answering the question: Given the training data and the available classes, what is the probability that an instance came from this class (instead of another of the available classes)? Equation 2 (Results), which defines the SRS, aims to ask a different question, which is simply: What is the likelihood of seeing an instance like this one, across all available classes?

In Figure 1 of this study, SWIF(r) was trained using a pi value of 0.5, an unbiased prior that gives both classes equal weight. In Figure 1A, data for each class was generated with a single attribute, drawn from N(−1, 0.2^2^) for class A and N(+1, 0.2^2^) for class B. In Figure 1C, we generated two attributes for each class instance. The values for the attributes were drawn randomly from a Gaussian distribution as before. However in this experiment, three different models were trained, each with their own parameters for the two Gaussian distributions (Table S2). 1000 instances were generated for each class to train SWIF(r). To create the testing set, 2000 instances were generated by drawing from the uniform distribution across the plane bounded by (−2, 2) in both x and y. SWIF(r) classification and the SRS were both calculated on each instance in the testing set, using the x and y coordinates as the attribute values, in order to observe SWIF(r) probabilities and SRS across this continuous range in attribute values. Data and code for this analysis are available on Github: https://github.com/ramachandran-lab/SRS_paper. SWIF(r), with the SRS included, can be installed according to instructions available on Github:https://github.com/ramachandran-lab/SWIFr.

### Unknown Class Example: Wheat seeds dataset

Data for this experiment was originally gathered by Charytanowicz et al. (2010). Three varieties of wheat: Kama (wheat 1), Rosa (wheat 2) and Canadian (wheat 3), were grown in experimental fields at the Institute of Agrophysics of the Polish Academy of Sciences in Lublin. 70 seeds from each variety were randomly selected, and visualized using a soft X-ray technique. Seven geometric parameters were measured: area, perimeter, compactness, length of kernel, width of kernel, asymmetry coefficient and length of kernel groove (Charytanowicz et al. 2010).

To construct training and testing sets, each set of 70 was randomly divided into two sets of 35, for training and testing respectively. We created three models, each trained on two types of wheat. Testing was then performed on all three types. Data and code for this analysis are available on Github: https://github.com/ramachandran-lab/SRS_paper.

### Systemic Mismatch between Training and Testing Set: UK Biobank elevated blood sugar

Phenotype data was downloaded from the UK Biobank under Application 22419, for 22 phenotypes used as attributes, and HBA1C. Attributes were as follows: BMI (Body Mass Index), MCV (Mean Corpuscular Volume), Platelet (count of Platelets in blood), DBP (Diastolic Blood Pressure), SBP (Systolic Blood Pressure), WBC (White Blood Cell count), RBC (Red Blood Cell count), Hemoglobin (Hemoglobin blood test), Hematocrit (hematocrit blood test), MCH (Mean Corpuscular Hemoglobin), MCHC (Mean Corpuscular Hemoglobin Concentration), Lymphocyte (Lymphocyte cell count), Monocyte (Monocyte cell count), Neutrophil (Neutrophil cell count), Eosinophil (Eosinophil cell count), Basophil (Basophil cell count), Urate (Uric acid concentration in blood), EGFR (Estimated Glomerular Filtration Rate), CRP (C-Reactive Protein concentration), Triglyceride (Triglyceride concentration), HDL (High-Density Lipoprotein test), LDL (Low-Density Lipoprotein test) and Cholesterol (Cholesterol test). For each phenotype, we centered the mean phenotype value at 0 and scaled the phenotype values to have a standard deviation of 1. We then selected a subset of those individuals for training each of two models. Within each model, individuals were divided on the basis of their HBA1C measurement, a blood test that estimates blood sugar levels. Individuals with an HBA1C reading of over 48 mmol/mol were sorted into the “elevated” cohort: readings of 48 mmol/mol or greater are typically considered diagnostic for diabetes. Individuals with an HBA1C reading of 42 mmol/mol or below were sorted into the “normal” cohort. Individuals with HBA1C readings of 42-48 mmol/mol, a range associated with prediabetes, were not included in the analysis.

#### Ancestry Mismatch Experiment

Individuals were first divided on the basis of ancestry, as in Smith et al. 2021, identifying 349,411 individuals of self-identified European descent, and 4,967 individuals of African descent. The latter of which were identified both by self-identification and by an ADMIXTURE analysis as described in (Smith et al. 2021). Applying the HBA1C filter described above resulted in 8,631 individuals in the European/elevated cohort, 268 individuals in the African/elevated cohort, 243,283 individuals in the European/normal cohort and 2,532 individuals in the African/normal cohort. For the purposes of training, 800 individuals were selected at random from each of the European/elevated and European/normal cohorts, creating a training set with 1600 European individuals. The model was then tested on 800 individuals from each cohort, with the exception of the African/elevated cohort which was tested on the 268 eligible individuals.

#### Sex Mismatch Experiment

The same 349,411 individuals of European descent described in the ancestry experiment were divided on the basis of sex, yielding 134,578 female individuals and 117,336 male individuals. Applying the HBA1C filter described above resulted in 131,432 individuals in the female/normal cohort, 3146 individuals in the female/elevated cohort, 111,851 individuals in the male/normal cohort and 5485 individuals in the male/elevated cohort. For the purposes of training, 800 individuals were selected at random from each of the male/normal and male/elevated cohorts, creating a training set with 1600 male individuals. The model was then tested on 800 individuals from each cohort.

### Missing Data: Balancing Selection/1000 Genomes dataset

To produce simulations of balancing selection and neutral evolution we used features of SLiM, pyslim and msprime to combine coalescent and forward simulation approaches (Haller et al. 2019). We simulated 1 Mb genomic regions with 500 simulations for each of the two conditions (balancing and neutral). Balancing simulations contained a single balanced mutation at the centerpoint of the 1 Mb genomic region. The mutation was introduced 100,000 generations ago, with a selection coefficient (s) of 0.01, and overdominance coefficient (h) varied according to a uniform distribution from 1.33 to 5. The simulation process used SLiM 3.3, including tree sequence recording, along with pyslim, tskit and msprime for recapitation and neutral mutation overlay (Haller et al. 2019). We simulated African, European, and Asian populations according to the demographic model developed by Gravel et al. 201 1. Following simulation, we sampled 200 individuals (400 haplotypes) from the African population for each selection scenario to generate training data. Testing data was generated in an identical manner, with 100 simulations performed for each selection scenario.

Cross-Population Extended Haplotype Homozygosity (XP-EHH) (Sabeti et al. 2007), Integrated Haplotype Score (iHS) (Voight et al. 2006), iHH12 (Garud et al. 2015; Torres, Szpiech, and Hernandez 2018) and nSL (Ferrer-Admetlla et al. 2014) were calculated using selscan (Szpiech and Hernandez 2014). These statistics use phased genomic data and were designed to detect signatures of recent or ongoing positive selection in genomes. Beta, a statistic built to detect balancing selection using ancestral/derived allele frequencies, was calculated using Betascan (Siewert and Voight 2017). For XP-EHH, which requires an outgroup population to calculate, the simulated European population served as the outgroup for the African population sampled. DDAF also requires an outgroup allele frequency to calculate. The average of the European and Asian allele frequencies was used.

For comparison with 1000 Genomes data, the YRI population was selected, and all of the above summary statistics were calculated on the complete set of YRI individuals. Where outgroup calculations were required, the CEU and CHB populations were used in the same manner as the European and Asian simulations, respectively. Simulated testing data for each scenario, as well as 1000 Genomes data was processed with SWIF(r), producing classification probabilities and SRS (shown in Figure 4A). Data and code for this analysis are available on Github: https://github.com/ramachandran-lab/SRS_paper.

## Acknowledgements

K. Ahlquist received funding from the NIH award 5T32GM007601 (to the department of Molecular Biology, Cell Biology and Biochemistry at Brown University), and from NIH grant R01GM118652 (to Sohini Ramachandran). This research was additionally supported by NIH grant R35GM139628 to Sohini Ramachandran. Lauren Sugden received funding from the Wimmer Family Foundation. The authors thank Samuel Pattillo Smith for assistance with UKBiobank data.

## References

1000 Genomes Project Consortium, Adam Auton, Lisa D. Brooks, Richard M. Durbin, Erik P. Garrison, Hyun Min Kang, Jan O. Korbel, et al. 2015. “A Global Reference for Human Genetic Variation.” Nature 526 (7571): 68–74.

Azodi, Christina B., Jiliang Tang, and Shin-Han Shiu. 2020. “Opening the Black Box: Interpretable Machine Learning for Geneticists.” Trends in Genetics: TIG 36 (6): 442–55.

Badgeley, Marcus A., John R. Zech, Luke Oakden-Rayner, Benjamin S. Glicksberg, Manway Liu, William Gale, Michael V. McConnell, Bethany Percha, Thomas M. Snyder, and Joel T. Dudley. 2019. “Deep Learning Predicts Hip Fracture Using Confounding Patient and Healthcare Variables.” NPJ Digital Medicine 2 (April): 31.

Bakker, Paul I. W. de, Gil McVean, Pardis C. Sabeti, Marcos M. Miretti, Todd Green, Jonathan Marchini, Xiayi Ke, et al. 2006. “A High-Resolution HLA and SNP Haplotype Map for Disease Association Studies in the Extended Human MHC.” Nature Genetics 38 (10): 1166–72.

Battey, C. J., Peter L. Ralph, and Andrew D. Kern. 2020a. “Space Is the Place: Effects of Continuous Spatial Structure on Analysis of Population Genetic Data.” Genetics 215 (1): 193–214.

Battey, C. J., Peter L. Ralph, and Andrew D. Kern. 2020b. “Predicting Geographic Location from Genetic Variation with Deep Neural Networks.” eLife 9 (June). https://doi.org/10.7554/eLife.54507.

Charytanowicz, Małgorzata, Jerzy Niewczas, Piotr Kulczycki, Piotr A. Kowalski, Szymon Łukasik, and Sławomir Żak. 2010. “Complete Gradient Clustering Algorithm for Features Analysis of X-Ray Images.” In Information Technologies in Biomedicine, 15–24. Springer Berlin Heidelberg.

Chen, Ruolan, Xiangrong Liu, Shuting Jin, Jiawei Lin, and Juan Liu. 2018. “Machine Learning for Drug-Target Interaction Prediction.” Molecules 23 (9). https://doi.org/10.3390/molecules23092208.

Eid, Fatma-Elzahraa, Haitham A. Elmarakeby, Yujia Alina Chan, Nadine Fornelos, Mahmoud ElHefnawi, Eliezer M. Van Allen, Lenwood S. Heath, and Kasper Lage. 2021. “Systematic Auditing Is Essential to Debiasing Machine Learning in Biology.” Communications Biology 4 (1): 183.

Ferrer-Admetlla, Anna, Mason Liang, Thorfinn Korneliussen, and Rasmus Nielsen. 2014. “On Detecting Incomplete Soft or Hard Selective Sweeps Using Haplotype Structure.” Molecular Biology and Evolution 31 (5): 1275–91.

Field, Yair, Evan A. Boyle, Natalie Telis, Ziyue Gao, Kyle J. Gaulton, David Golan, Loic Yengo, et al. 2016. “Detection of Human Adaptation during the Past 2000 Years.” Science, November. https://doi.org/10.1126/science.aag0776.

Freitas, Alex A. 2014. “Comprehensible Classification Models: A Position Paper.” SIGKDD Explor. Newsl. 15 (1): 1–10.

Garud, Nandita R., Philipp W. Messer, Erkan O. Buzbas, and Dmitri A. Petrov. 2015. “Recent Selective Sweeps in North American Drosophila Melanogaster Show Signatures of Soft Sweeps.” PLoS Genetics 11 (2): e1005004.

Gilpin, William, Yitong Huang, and Daniel B. Forger. 2020. “Learning Dynamics from Large Biological Datasets: Machine Learning Meets Systems Biology.” Current Opinion in Systems Biology. https://www.sciencedirect.com/science/article/pii/S2452310020300147.

Gravel, Simon, Brenna M. Henn, Ryan N. Gutenkunst, Amit R. Indap, Gabor T. Marth, AndrewG. Clark, Fuli Yu, Richard A. Gibbs, 1000 Genomes Project, and Carlos D. Bustamante. 2011. “Demographic History and Rare Allele Sharing among Human Populations.” Proceedings of the National Academy of Sciences of the United States of America 108 (29): 11983–88.

Greener, Joe G., Shaun M. Kandathil, Lewis Moffat, and David T. Jones. 2022. “A Guide to Machine Learning for Biologists.” Nature Reviews. Molecular Cell Biology 23 (1): 40–55.

Haller, Benjamin C., Jared Galloway, Jerome Kelleher, Philipp W. Messer, and Peter L. Ralph. 2019. “Tree-Sequence Recording in SLiM Opens New Horizons for Forward-Time Simulation of Whole Genomes.” Molecular Ecology Resources 19 (2): 552–66.

Han, Yi, Juze Yang, Xinyi Qian, Wei-Chung Cheng, Shu-Hsuan Liu, Xing Hua, Liyuan Zhou, et al. 2019. “DriverML: A Machine Learning Algorithm for Identifying Driver Genes in Cancer Sequencing Studies.” Nucleic Acids Research 47 (8): e45.

Heil, Benjamin J., Michael M. Hoffman, Florian Markowetz, Su-In Lee, Casey S. Greene, and Stephanie C. Hicks. 2021. “Reproducibility Standards for Machine Learning in the Life Sciences.” Nature Methods 18 (10): 1132–35.

Jones, David T. 2019. “Setting the Standards for Machine Learning in Biology.” Nature Reviews.Molecular Cell Biology 20 (11): 659–60.

Lenz, Tobias L., Victor Spirin, Daniel M. Jordan, and Shamil R. Sunyaev. 2016. “Excess of Deleterious Mutations around HLA Genes Reveals Evolutionary Cost of Balancing Selection.” Molecular Biology and Evolution 33 (10): 2555–64.

Lewontin, R. C., and J. Krakauer. 1973. “Distribution of Gene Frequency as a Test of the Theory of the Selective Neutrality of Polymorphisms.” Genetics 74 (1): 175–95.

Linardatos, Pantelis, Vasilis Papastefanopoulos, and Sotiris Kotsiantis. 2020. “Explainable AI: A Review of Machine Learning Interpretability Methods.” Entropy 23 (1). https://doi.org/10.3390/e23010018.

Lipton, Zachary C. 2018. “The Mythos of Model Interpretability: In Machine Learning, the Concept of Interpretability Is Both Important and Slippery.” Queueing Systems. Theory and Applications 16 (3): 31–57.

Lürig, Moritz, Seth Donoughe, Erik Svensson, Arthur Porto, and Masahito Tsuboi. 2021. “Computer Vision, Machine Learning, and the Promise of Phenomics in Ecology and Evolutionary Biology.” https://doi.org/10.32942/osf.io/98cuw.

Murphy, Kevin P. 2012. Machine Learning: A Probabilistic Perspective. MIT Press.

Pavlidis, Pavlos, Jeffrey D. Jensen, and Wolfgang Stephan. 2010. “Searching for Footprints of Positive Selection in Whole-Genome SNP Data from Nonequilibrium Populations.” Genetics 185 (3): 907–22.

Poplin, Ryan, Avinash V. Varadarajan, Katy Blumer, Yun Liu, Michael V. McConnell, Greg S. Corrado, Lily Peng, and Dale R. Webster. 2018. “Prediction of Cardiovascular Risk Factors from Retinal Fundus Photographs via Deep Learning.” Nature Biomedical Engineering 2 (3): 158–64.

Rudin, Cynthia. 2019. “Stop Explaining Black Box Machine Learning Models for High Stakes Decisions and Use Interpretable Models Instead.” Nature Machine Intelligence. https://doi.org/10.1038/s42256-019-0048-x.

Rudin, Cynthia, Chaofan Chen, Zhi Chen, Haiyang Huang, Lesia Semenova, and Chudi Zhong. 2022. “Interpretable Machine Learning: Fundamental Principles and 10 Grand Challenges.” Statistics Surveys 16 (none): 1–85.

Sabeti, Pardis C., Patrick Varilly, Ben Fry, Jason Lohmueller, Elizabeth Hostetter, Chris Cotsapas, Xiaohui Xie, et al. 2007. “Genome-Wide Detection and Characterization of Positive Selection in Human Populations.” Nature 449 (7164): 913–18.

Sankararaman, Sriram, Swapan Mallick, Michael Dannemann, Kay Prüfer, Janet Kelso, Svante Pääbo, Nick Patterson, and David Reich. 2014. “The Genomic Landscape of Neanderthal Ancestry in Present-Day Humans.” Nature 507 (7492): 354–57.

Schrider, Daniel R., and Andrew D. Kern. 2016. “S/HIC: Robust Identification of Soft and Hard Sweeps Using Machine Learning.” PLoS Genetics 12 (3): e1005928.

Siewert, Katherine M., and Benjamin F. Voight. 2017. “Detecting Long-Term Balancing Selection Using Allele Frequency Correlation.” Molecular Biology and Evolution 34 (11): 2996–3005.

Smith, Samuel Pattillo, Sahar Shahamatdar, Wei Cheng, Selena Zhang, Joseph Paik, Misa Graff, Christopher Haiman, et al. 2021. “Enrichment Analyses Identify Shared Associations for 25 Quantitative Traits in over 600,000 Individuals from Seven Diverse Ancestries.” bioRxiv. https://doi.org/10.1101/2021.04.20.440612.

Sugden, Lauren Alpert, Elizabeth G. Atkinson, Annie P. Fischer, Stephen Rong, Brenna M. Henn, and Sohini Ramachandran. 2018. “Localization of Adaptive Variants in Human Genomes Using Averaged One-Dependence Estimation.” Nature Communications 9 (1): 703.

Szpiech, Zachary A., and Ryan D. Hernandez. 2014. “Selscan: An Efficient Multithreaded Program to Perform EHH-Based Scans for Positive Selection.” Molecular Biology and Evolution 31 (10): 2824–27.

Tajima, F. 1989. “Statistical Method for Testing the Neutral Mutation Hypothesis by DNA Polymorphism.” Genetics 123 (3): 585–95.

Torres, Raul, Zachary A. Szpiech, and Ryan D. Hernandez. 2018. “Human Demographic History Has Amplified the Effects of Background Selection across the Genome.” PLoS Genetics 14 (6): e1007387.

Villanea, Fernando A., and Joshua G. Schraiber. 2018. “Multiple Episodes of Interbreeding between Neanderthal and Modern Humans.” Nature Ecology & Evolution 3 (1): 39–44.

Villoutreix, Paul. 2021. “What Machine Learning Can Do for Developmental Biology.” Development 148 (1). https://doi.org/10.1242/dev.188474.

Voight, Benjamin F., Sridhar Kudaravalli, Xiaoquan Wen, and Jonathan K. Pritchard. 2006. “A Map of Recent Positive Selection in the Human Genome.” PLoS Biology 4 (3): e72.

Walsh, Ian, Dmytro Fishman, Dario Garcia-Gasulla, Tiina Titma, Gianluca Pollastri, ELIXIR Machine Learning Focus Group, Jennifer Harrow, Fotis E. Psomopoulos, and Silvio C. E. Tosatto. 2021. “DOME: Recommendations for Supervised Machine Learning Validation in Biology.” Nature Methods 18 (10): 1122–27.

Weir, Bruce S. 1990. Genetic Data Analysis. Methods for Discrete Population Genetic Data. Sinauer Associates, Inc. Publishers.

Wojtusiak, Janusz. 2021. “Reproducibility, Transparency and Evaluation of Machine Learning in Health Applications.” Proceedings of the 14th International Joint Conference on Biomedical Engineering Systems and Technologies. https://doi.org/10.5220/0010348306850692.

Zangooei, Mohammad Hossein, and Jafar Habibi. 2017. “Hybrid Multiscale Modeling and Prediction of Cancer Cell Behavior.” PloS One 12 (8): e0183810.

Zitnik, Marinka, Francis Nguyen, Bo Wang, Jure Leskovec, Anna Goldenberg, and Michael M. Hoffman. 2019. “Machine Learning for Integrating Data in Biology and Medicine: Principles, Practice, and Opportunities.” An International Journal on Information Fusion 50 (October): 71–91.

