## Supplemental Tables for "Enabling interpretable machine learning for biological data with reliability scores"

### Supplementary Material

**Table S1. Average SRS values and p-values for UKB cohort comparisons**

| Cohort | Mean SRS* | p-value[†] |
|---|---|---|
| Male Elevated | -10.04 | 1.53e-4 |
| Female Elevated | -11.14 | |
| Male Normal | -8.57 | 5.11e-34 |
| Female Normal | -10.47 | |
| European Elevated | -10.47 | 6.04e-6 |
| African Elevated | -11.90 | |
| European Normal | -9.22 | 1.50e-28 |
| African Normal | -11.10 | |

*SRS truncated to second decimal place

[†]calculated using a t-test: scipy.stats.ttest_ind(cohort_a_SRS, cohort_b_SRS)

**Table S2. Gaussian distributions by class and model for Figure 1C**

| | Model 1 | Model 2 | Model 3 |
|---|---|---|---|
| Class 1 | Mean = -1,-1<br>Covariance Array =<br>np.array([[0.1, 0.085],<br>[0.085, 0.1]]) | Mean = -0.5,0<br>Covariance Array =<br>np.array([[0.25,0.],[0.,0.25]<br>]) | Mean = 0.25, 0.25<br>Covariance Array =<br>np.array([[-0.25, 0.2], [0.2,<br>-0.25]]) |
| Class 2 | Mean = 1,1<br>Covariance Array =<br>np.array([[0.1, 0.085],<br>[0.085, 0.1]]) | Mean = 0.5,0<br>Covariance Array =<br>np.array([[0.25,0.],[0.,0.25]<br>]) | Mean = -0.5, -0.5<br>Covariance Array =<br>np.array([[0.3,0.],[0.,0.3]]) |
